# Interleukin 4 controls the role of macrophages in pulmonary metastatic tumor cell seeding and growth

**DOI:** 10.1101/2021.12.28.474388

**Authors:** Carolina Rodriguez-Tirado, David Entenberg, Jiufeng Li, Bin-Zhi Qian, John S. Condeelis, Jeffrey W Pollard

## Abstract

Metastasis is the systemic manifestation of cancer and the main cause of death from breast cancer. In mouse models of lung metastases, recruitment of classical monocytes from blood to the lung and their differentiation to metastasis-associated macrophages (MAMs) facilitate cancer cell extravasation, survival, and growth. Ablation of MAMs or their monocytic progenitors inhibits metastasis. We hypothesized that factors controlling macrophage polarization modulate tumor cell extravasation in the lung. We evaluated whether signaling by Th1 or Th2 cytokines in macrophages affected trans-endothelial migration of tumor cells in vitro. Interferon γ and LPS inhibited macrophage-dependent tumor cell extravasation while the Th2 cytokine interleukin-4 (IL4) enhanced this process. We demonstrated that IL4 receptor (*IL4rα)* null mice develop fewer and smaller lung metastasis. Adoptive transfer of wild type monocytes to *IL4rα* deficient mice rescued this phenotype. IL4 signaling in macrophages controls the expression of the chemokine receptor CXCR2, necessary for IL4-mediated tumor cell extravasation in vitro. Furthermore, IL4 signaling in macrophages transcriptionally regulates several other genes already causally associated with lung metastasis including CCL2, CSF1, CCR1, HGF and FLT1. The central role for IL4 signaling in MAMs was confirmed by high-resolution intravital imaging of the lung in mice at the time of metastatic seeding, which showed reduced physical interaction between tumor cells and *IL4rα*-deficient macrophages. This interaction enhances tumor cell survival. These data indicate that IL4 signaling in monocytes and macrophages is key during seeding and growth of breast metastasis in the lung as it regulates pro-tumoral paracrine signaling between cancer cells and macrophages.

## Introduction

Breast cancer is the single most commonly diagnosed cancer among women worldwide [1, 2] and it is estimated to be responsible for the death of over 41,000 women in the US alone during 2019 [3]. The vast majority of cancer-related deaths, including those from breast cancer, are a result of metastatic disease and its associated complications. The fact that survival of patients with metastatic disease has hardly improved for two decades highlights that early detection and effective treatment of established metastases is an essential clinical need.

Most experimental studies as well as sequencing studies of cancer mutations have focused upon genetic and epigenetic changes that occur in cancer cells to enable metastases to prosper and occupy particular anatomical sites. However, it has become apparent, both in primary tumors and in the seeding and expansion stages of metastatic tumors, that the environment that tumors create determines their survival and growth. This tumor microenvironment (TME), in both humans and mice, is populated by many cell types. Of these, immune cells, and macrophages in particular, are highly abundant. In breast cancer, the importance of macrophages in disease progression was initially shown via genetic or pharmacological depletion in animal models, which suppressed lung metastasis [4, 5]. Since then, similar studies in mouse models have also been performed in many other cancer types including glioblastoma and pancreatic cancer [6, 7]. The importance of macrophages in human cancers is also supported by numerous (though not all) clinical correlative studies that have shown a strong correlation between high macrophage infiltration and poor prognosis [8-11]. Recently individual macrophage transcriptional signals have also shown to be independent prognostic indicators of poor disease specific survival [12].

While an abundant body of information is available regarding the pro-tumoral role of macrophages at the primary site, it is still unclear whether the same mechanisms control their recruitment and activities at sites of metastases. Our previous work showed that in pulmonary foci from mouse models of metastatic breast cancer, expression of the VEGF receptor type 1, FMS-like tyrosine kinase 1 (FLT1), allows the identification of a particular population of macrophages (herein called metastasis-associated macrophages (MAMs)), that differ phenotypically and functionally from both tumor associated-macrophages (TAMs) at the primary site, and myeloid resident cells in the lung [13-15]. We showed that MAMs differentiate from circulating classical Ly6C+ CCR2+ monocytes (CMs) that are recruited to the metastases by tumor cell-derived CCL2 [15, 16]. Furthermore, once recruited, they establish direct contact with cancer cells and promote their extravasation and survival [5]. Recruitment of CMs by CCL2 is essential for metastatic progression as blocking CCL2 reduces seeding and growth of pulmonary metastasis [16]. These CMs differentiate into a MAM intermediate precursor (MAMPC) state (with properties of myeloid suppressor cells, MDSC) and then fully differentiate into MAMs [15]. MAMPCs respond to CCL2 by inducing the expression of CCL3 (which acts in an autocrine manner through its receptor CCR1). This promotes MAMPC retention at the metastatic site by enhancing their physical interaction with tumor cells through VCAM1. This interaction delivers a survival signal that further favors tumor cell extravasation and disease progression [17, 18].

The TME in primary tumors is thought to be skewed to a Th2 type (a pro-tumoral immune response) rather than an inflammatory Th1 one (regarded as anti-tumoral). Supporting this hypothesis, the Th2 regulators, Interleukin-4 (IL4) and 13 (IL13), as well as IL10, have been demonstrated to play important roles during primary tumor progression in mouse models of cancer [19-21]. Similarly, IL10 inhibits anti-tumoral cytotoxic T cell responses thereby promoting tumor progression. However, in this case, the mechanism appears to be via effects on dendritic cell recruitment and function, and not through TAMs [19]. In humans, high expression levels of IL4 are observed in breast tumors, which is associated with a Th2-skewed profile of tumor-infiltrating lymphocytes [22, 23].

IL4 signaling can be through either a type I receptor where the alpha chain binds to a common gamma chain (γc) or a type II receptor, where the alpha chain interacts with IL13Ra1 to form a dimer. The type II receptor allows IL13 signaling. In the TME as well as IL4rα expression on many myeloid cells, tumor cell lines derived from primary human breast [24, 25] and lung carcinomas [26] have also been observed to have increased levels of IL4 receptors. Together these data suggest that IL4 signaling is important for the *in vivo* control of tumor progression by targeting both tumor cells and innate immune cells in the TME.

Though much is known about primary tumors, less is known about mechanisms that regulate immunity in the metastatic site. In mouse models of bone metastasis, IL4 has also been shown to polarize bone MAMs (BoMAMs)to promote tumor growth [27]. In the lung however, it is still unknown whether IL4 signaling participates in metastatic seeding and growth of tumor cells, or whether its role is mainly restricted to the primary tumor and bone metastases. Thus, we sought to determine whether IL4 signaling is relevant for macrophage function during lung metastatic seeding. In an experimental metastasis model that eliminates the contribution of the primary tumor, we determined that IL4 signaling in macrophages facilitates seeding of mammary tumor cells to the lung, thus suggesting a specific role for this cytokine at the metastatic site. We identified CXCR2 as an important downstream factor from IL4Rα that promotes extravasation of cancer cells. Furthermore, IL4 increases expression of genes shown to be essential for metastatic seeding and growth including *Ccr1, Flt1, Ccl3, Hgf, Csf1*. Importantly, high-resolution *in vivo* imaging of IL4rα null mice at early stages of lung colonization demonstrates that IL4rα null macrophages show reduced levels of physical interaction with tumor cells, which provides a mechanistic explanation for the observed reduced seeding and growth of mammary tumors in the lung.

## Results

### Interleukin 4 signaling in macrophages enhances cancer cell trans-endothelial migration

In vitro assays that mimic extravasating tumor cell transendothelial migration (eTEM) at distant sites have been useful for identifying the signaling pathways by which macrophages promote extravasation *in vivo* activities with high fidelity (**Figure 1A**) [5, 16, 28]. Therefore, we utilized this eTEM assay to evaluate whether Th2 or Th1 polarization plays a role in regulating macrophage promotion of tumor cell transendothelial cell migration. In order to study mammary cancer metastasis, the transendothelial migration of the tumor cell lines Met-1 (derived from a PyMT mammary tumor) and E0771-LG (derived from a medullary mammary cancer selected for high metastatic potential) in response to bone marrow derived macrophages (BMDMs) was analyzed as previously described [28]. To confirm the presence of a tight endothelial barrier in the assay, we measured the permeability of the endothelial layer using both 70 and 10 KDa dextrans as well as the electrical resistance across the barrier (**Supplemental Figure 1A, B**), as previously described [28]. After this eTEM assay optimization, IL4Rα signaling was analyzed. First, the effect of an exogenous source of IL4 on tumor cell trans-endothelial migration events *in vitro* in the presence of macrophages was measured. Wild type (WT) BMDMs were pre-incubated with recombinant IL4 and placed into the assay. This IL4 treatment induced a significant increase in the number of E0771-LG (n=3, <0.005) (**Figure 1B)** and Met-1 (not shown, n=3, p-value <0.0001) cells that migrated across the endothelial barrier compared to untreated controls. This response was specific to IL4 signaling, as IL4 stimulation of BMDMs isolated from *IL4Rα*^*-/-*^ mice did not enhance tumor cell transendothelial migration (**Figure 1B**; n=3, p-value >0.05). However, in contrast to this enhancement by IL4, there was no significant difference in the number of either Met-1 or E0771-LG tumor cells that transmigrated the endothelial barrier in response to *IL4Rα*^*+/-*^ or *IL4Rα*^*-/-*^ BMDMs compared to WT BMDMs (Met-1 cells, n=5, p-value_**(*IL4Rα+/-*)**_=0.59 and p-value _**(*IL4Rα-/-*)**_=0.15; E0771-LG cells, n=4, p-value_**(*IL4Rα+/-*)**_=0.88 and p-value_**(*IL4Rα-/-*)**_=0.98) (**Supplemental Figure 1C**). Similarly, BMDMs isolated from mice lacking the γC receptor, a sub-unit that is required for the formation of the type I IL4R complex, did not alter the transendothelial migration of Met-1 cells (Met-1 cells, n=3, p-value=0.14) (**Figure 1C**). Thus, IL4 signaling is not required for macrophage enhanced tumor cell transendothelial migration in this assay but IL4 potentiated the macrophage activity. To evaluate whether IL13 (a cytokine sharing the type II IL4R) mediates a similar response, BMDMs were stimulated with recombinant IL13 and the tumor cell transendothelial migration measured. In contrast to IL4, there was no change in the number of transmigrated tumor cells after pre-stimulation of BMDMs with IL13 (Met-1 cells, n=3, p-value=0.33) (**Figure 1D**).

**Figure 1.**
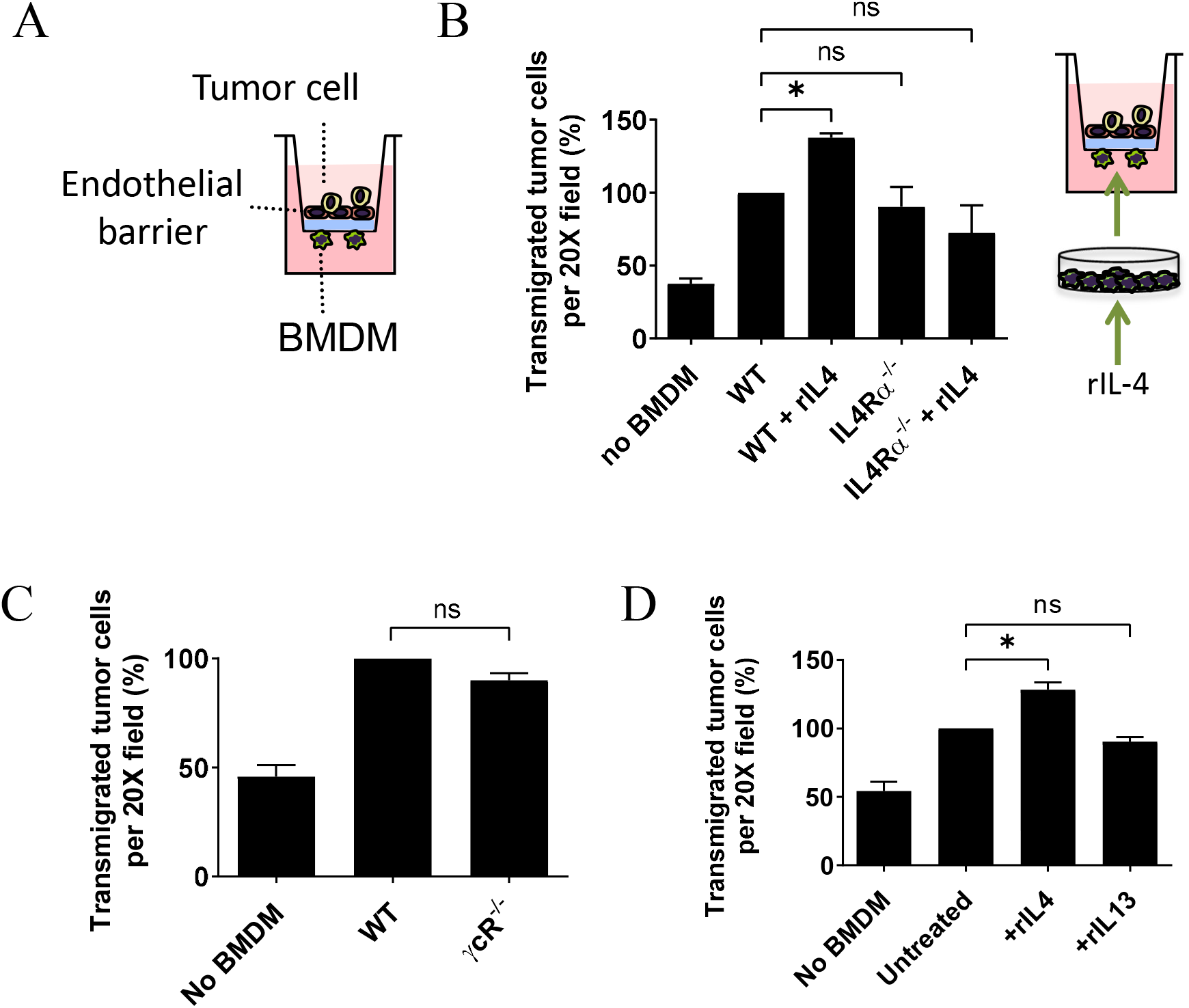
IL4 promotes macrophage-mediated tumor cell trans-endothelial migration. **(A)** Schematics of the transendothelial migration assay modeling extravasation (eTEM). An endothelial monolayer (red) sits over a Matrigel-coated transwell. Endothelial barrier is exposed to tumor cells on the apical domain. Macrophages are added to the bottom side of the transwell insert (closer to basal domain of the endothelial barrier). Tumor cells that migrate through the endothelial layer and matrigel are quantified once present in the bottom side of the transwell insert. (**B**) Transendothelial migration of E0771-LG cells in response to WT or *IL4Rα* null BMDMs pre-incubated with vehicle or rIL4. ANOVA, n=3, * p-value<0.05. **(C)** Transendothelial migration of Met-1 cells in presence of BMDM deficient for the IL4R co-receptor γCR. Mann Whitney test, n=3, p-value>0.1. **(D)** Trans-endothelial migration of Met-1 cells in response to WT BMDMs pre-incubated with recombinant IL4 or IL13 for 24 h. Mann-Whitney test, n=3, * p-value<0.01. Data is presented as the mean normalized to WT or untreated control +/-SD. Each experiment was performed in duplicate or triplicate, and each n corresponds to an independent experiment performed with BMDM from independent mice.

Macrophages in the primary tumor TME can synthesize IL10, another Th2 cytokine [19]. Thus, to study IL10 signaling in macrophage promotion of tumor cell transendothelial migration, eTEM assays were used with BMDMs with and without a neutralizing antibody to IL10 receptor. IL10 receptor neutralizing antibody was applied either in the bottom or both chambers in the absence of exogenous IL10. This neutralizing antibody had no effect on the promotion by BMMs of Met-1 transendothelial migration (**Supplemental Figure 1D**). To further investigate IL10 signaling, BMMs were pre-incubated with IL10 or not and compared in the eTEM assay. However, this IL10 treatment made no significant difference in their activity compared to untreated BMMs in the eTEM assay (**Supplemental Figure 1E**). IL10 signals via signal transducer and activator of transcription (STAT3), the central transcriptional effector in macrophages. *Stat3* null mutant mice are not viable. Thus, to study the dependence on STAT3 in the eTEM assay, BMMs were derived from a homozygous floxed allele for *Stat3* crossed onto a myeloid restricted *Csf1* receptor (*Csf1R)* improved cre recombinase strain [29, 30]. BMMs when *Stat3* was deleted by the presence of the *Csf1*.*icre* were not significantly different in their promotion activity than from those derived from either WT or *Stat3*^*fl/fl*^ mice (**Supplemental Figure 1F**). We can thus conclude that IL10-STAT3 signaling does not promote or inhibit macrophage stimulation of tumor cell transendothelial migration *in vitro*.

To assess the role of Th1 cytokines and stimulators of macrophage activation in the eTEM assay BMMs were treated with INFγ, the TLR4 agonist lipopolysaccharide (LPS), or both together. In all cases, these treatments completely abolished the macrophage stimulation of tumor cell transendothelial cell migration to the basal level achieved by tumor cells alone (**Supplemental Figure 1G**).

Together, these results show that WT BMMs promote tumor cell trans-endothelial migration *in vitro* while IL4Rα signaling from IL4 (but not other Th2 or Th1 cytokines) in macrophages enhances their activity in this process. These *in vitro* data lead us to hypothesize that the lung metastatic TME is aided by Th2 polarization at the seeding step.

### IL4Rα is important for efficient lung metastasis development

To test this hypothesis that IL4 drives metastasis *in vivo*, an experimental metastasis assay using tail vein injections of syngeneic E0771-LG cells was employed to compare disease load in *IL4Rα* null mutant compared to WT mice, both on a BL6 background. Lungs were collected 11 days after injection and processed for metastatic burden quantification (**Figure 2A**). Rigorous stereological quantification of tumor burden in the lungs [5] showed that *IL4Rα* null mice developed fewer foci (**Figure 2B**) (n=10, p-value <0.005), with decreased size (**Figure 2C**) compared to WT mice (n=10, p-value <0.05), indicative of reduced seeding and growth of cancer cells, respectively. The metastatic index, a relationship between tumor volume compared to lung volume that indicates total metastatic load, was also decreased in *IL4Rα*^*-/-*^ mice (n=10, p-value<0.05) (**Figure 2D**). Although there was a trend towards a reduced foci number, foci size and metastatic index in heterozygous mice (*IL4Rα*^*+/-*^) compared to WT mice this reduction did not reach statistical significance when comparing WT and *IL4Rα*^*+/-*^ mice or *IL4Rα*^*+/-*^ and *IL4Rα*^*-/-*^ mice (**Figure 2B&C&D**). These results indicate that IL4 signaling through *IL4Rα* is important for tumor cell seeding and growth *in vivo*.

**Figure 2.**
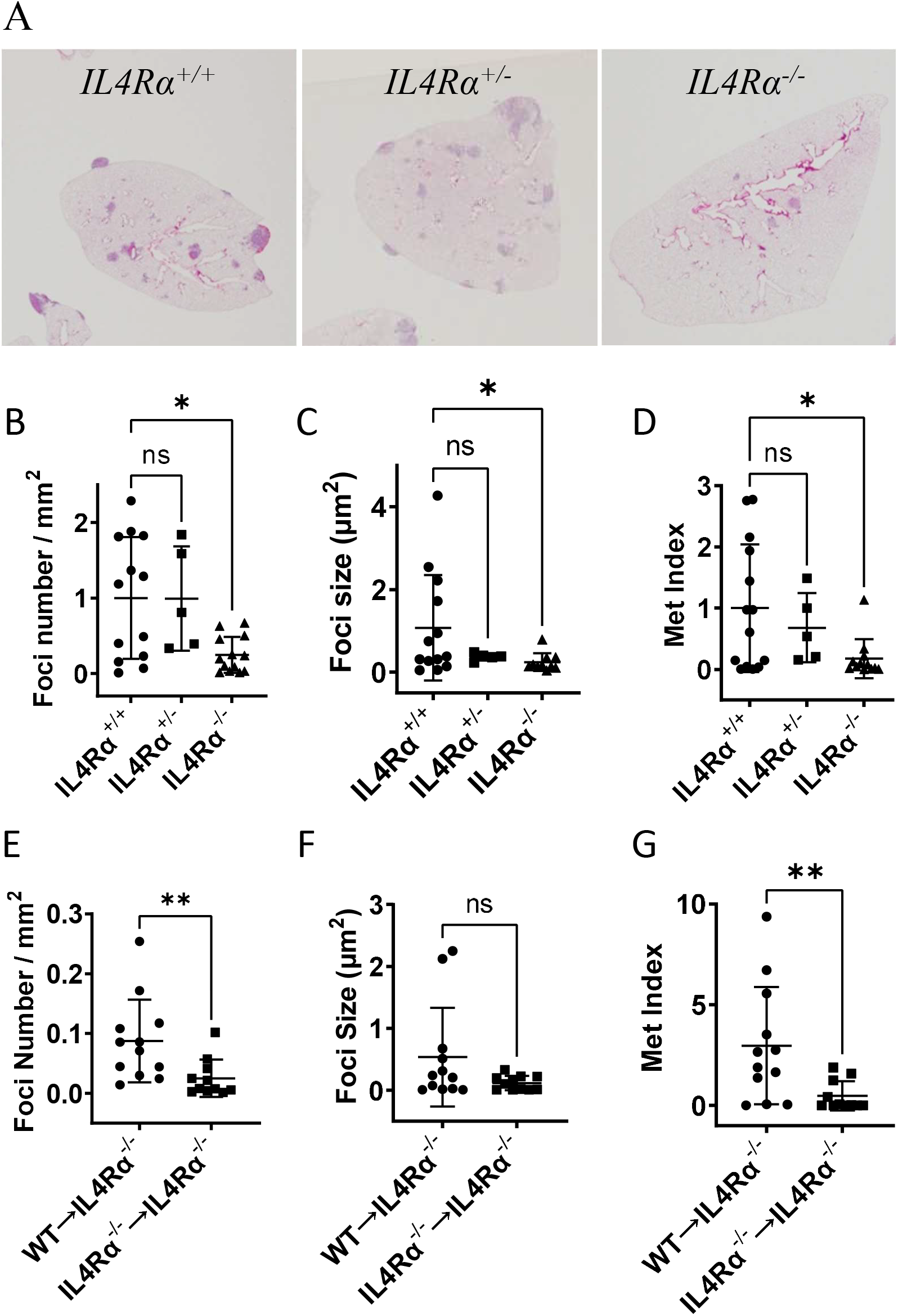
IL4 signaling is important for pulmonary metastatic colonization. **(A)** Hematoxylin & eosin-stained sections of the lungs from WT, *IL4Rα*^+/-^ and *IL4Rα*^-/-^ mice intravenously injected with syngeneic E0771-LG cells 11 days post-injection. **(B)** Stereological quantification of foci number in WT, *IL4Rα*^+/-^ and *IL4Rα*^-/-^ mice 11 days after intravenous injection of E0771-LG cells by counting the number of all foci every 100 μm interval in a total of six sections. Graphs correspond to 5 to 13 mice per group with bars representing average +/-SD. Kruskal-Wallis test, *p-value<0.05. **(C)** Stereological quantification of foci size in WT, *IL4Rα*^+/-^ and *IL4Rα*^-/-^ mice 11 days after intravenous injection of E0771-LG cells, calculated after measuring the area of all foci every 100 μm interval in a total of six sections. Graphs correspond to 5 to 13 mice per group with bars representing average +/-SD. Kruskal-Wallis test, * p-value<0.05 **(D)** Stereological quantification of the metastatic index, the ratio of total foci area by lung area, in WT, *IL4Rα*^+/-^ and *IL4Rα*^-/-^ mice 11 days after intravenous injection of E0771-LG cells Graphs correspond to 5 to 13 mice per group with bars representing average +/-SD. Kruskal-Wallis test, * p-value<0.05. **(E)** Stereological quantification of foci number in *IL4Rα*^*-/-*^ mice adoptively transferred with WT (WT→*IL4Rα*^-/-^) or *IL4Rα*^-/-^ (*IL4Rα*^-/-^→ *IL4Rα*^-/-^) monocytes immediate after E0771-LG cell injection. Graphs correspond to 11 to 12 mice per group with bars representing average +/-SD. Mann-Whitney test, n>11/group, ** p-value<0.005. **(F)** Stereological quantification of foci size in *IL4Rα*^*-/-*^ mice adoptively transferred with WT (WT→*IL4Rα*^-/-^) or *IL4Rα*^-/-^ (*IL4Rα*^-/-^→ *IL4Rα*^-/-^) monocytes immediate after E0771-LG cell injection. Graphs correspond to 11 to 12 mice per group with bars representing average +/-SD. Mann-Whitney test, n>11/group, p-value>0.05. **(G)** Stereological quantification of metastatic index in *IL4Rα*^*-/-*^ mice adoptively transferred with WT (WT→*IL4Rα*^-/-^) or *IL4Rα*^-/-^ (*IL4Rα*^-/-^→ *IL4Rα*^-/-^) monocytes immediate after E0771-LG cell injection. Graphs correspond to 11 to 12 mice per group with bars representing average +/-SD. Mann-Whitney test, n>11/group, ** p-value<0.005.

The null mutation in *IL4Rα* mice affects signaling in all cell types where it is expressed. Thus, to determine if IL4 signaling in monocytes is responsible for the stimulation of lung metastasis, classical monocytes (CM) were isolated from WT or *IL4Rα*^*-/-*^ mice and adoptively transferred into *IL4Rα* null mice (WT→*IL4Rα*^*-/-*^; *IL4Rα*^*-/-*^→*IL4Rα*^*-/-*^) and the effect on metastasis evaluated. We observed that WT CMs rescued the foci number (**Figure 2E**) and total metastatic burden (metastatic index) (**Figure 2G**) in *IL4Rα*^*-/-*^ null mice while *IL4Rα*^*-/-*^ CMs did not (n>11, p-value 0.012). In these experiments, focal size (**Figure 2F**) was not significantly restored, probably due to the short life span of the adoptively transferred CMs that were only transferred once at the beginning of the assay. Nevertheless, these gain-of-function over loss-of-function results show that a deficiency in IL4 signaling in monocytes/macrophages is responsible for the reduced metastatic burden in *IL4Rα* mice.

Initial seeding of metastatic cells is enhanced by classical monocytes [16], thus we looked to determine if their number was reduced in the *IL4Rα*^*-/-*^ mice. No significant difference in the number of circulating classical (CM: CD45^+^ CD11b^+^ F4/80^+^ Ly6G^-^ Ly6C^hi^) or non-classical patrolling monocytes (PM: CD45^+^ CD11b^+^ F4/80^+^ Ly6G^-^ Ly6C^-^) from the total CD45^+^ CD11b^+^ F4/80^+^ cells was observed between WT and *IL4Rα*^*-/-*^ mice (n>9, p-value>0.15) (**Supplementary Figure 2**). Thus, the absence of *IL4Rα* does not affect the number of circulating monocytes available to be recruited by tumor cells to the lungs and, consequently, the reduced tumor burden in the lung of *IL4Rα*^*-/-*^ mice is not due to a reduction in the availability of circulating monocytes but their function.

### IL4 Signaling stabilize tumor cell-macrophage contact in vivo

Previous data indicated that CCL3/CCR1 signaling regulates tumor cell macrophage interactions that, once established, deliver a survival signal to the tumor cells through engagement of αV-integrin expressed on MAMs and VECAM1 on the tumor cell. This enhances metastatic cell seeding capacity as inhibition of this signaling pathway or of monocyte recruitment inhibited metastases [16-18]. Here, we used a novel multiphoton intravital imaging of the lung with a vacuum stabilized imaging window [31, 32] coupled with quantitative measurements to understand the dynamics of monocyte recruitment and differentiation (into MAMs), as well as their interaction with tumor cells during tumor cell lodging and seeding. In previous reports using MacBlue mice in which monocytes and macrophages are labelled with CFP [31, 33], we could phenotypically identify monocytes as small CFP+ cells in the circulation and differentiate them from tissue resident macrophages, which showed static behavior and extravascular location [32]. We previously showed that even single-cell stage tumor foci were characteristically associated with macrophages and that there was a tight physical interaction with macrophages without signs of tumor cell death [31]. In the current experiments, metrics were developed based upon cell tracking and were used to quantify the time of tumor cell-monocyte and tumor cell-macrophage interactions in WT and *IL4rα* mutant mice. Upon arrival in the lung of WT mice, intravascular tumor cells had limited interactions with circulating monocytes, lasting an average of 14±2 min (n=50). In *IL4rα* null mutant mice, the initial interactions between tumor cells and monocytes were similar to that observed in WT mice, with an average interaction time of 20.3±3 min (n=46, p>0.05) (**Figure 3A, D; Suppemental Movie 1**). However, in WT mice, as soon as tumor cells had extravasated, they displayed stable physical contact with macrophages that lasted an average of 195±30 min (n=25), significantly longer than tumor cell-monocyte interactions (p<0.0001) **(Figure 3B, D; Supplemental movie 2)**. Importantly, this time of interaction of tumor cells with *IL4rα*^-/-^ macrophages was significantly shorter, with an average of 84.13±12 min (n=43, p<0.0001) **(Figure 3C, D; Supplemental movie 3)**. Thus, IL4 signaling maintains the stability of MAM-tumor cell interactions but not their initial interactions.

**Figure 3.**
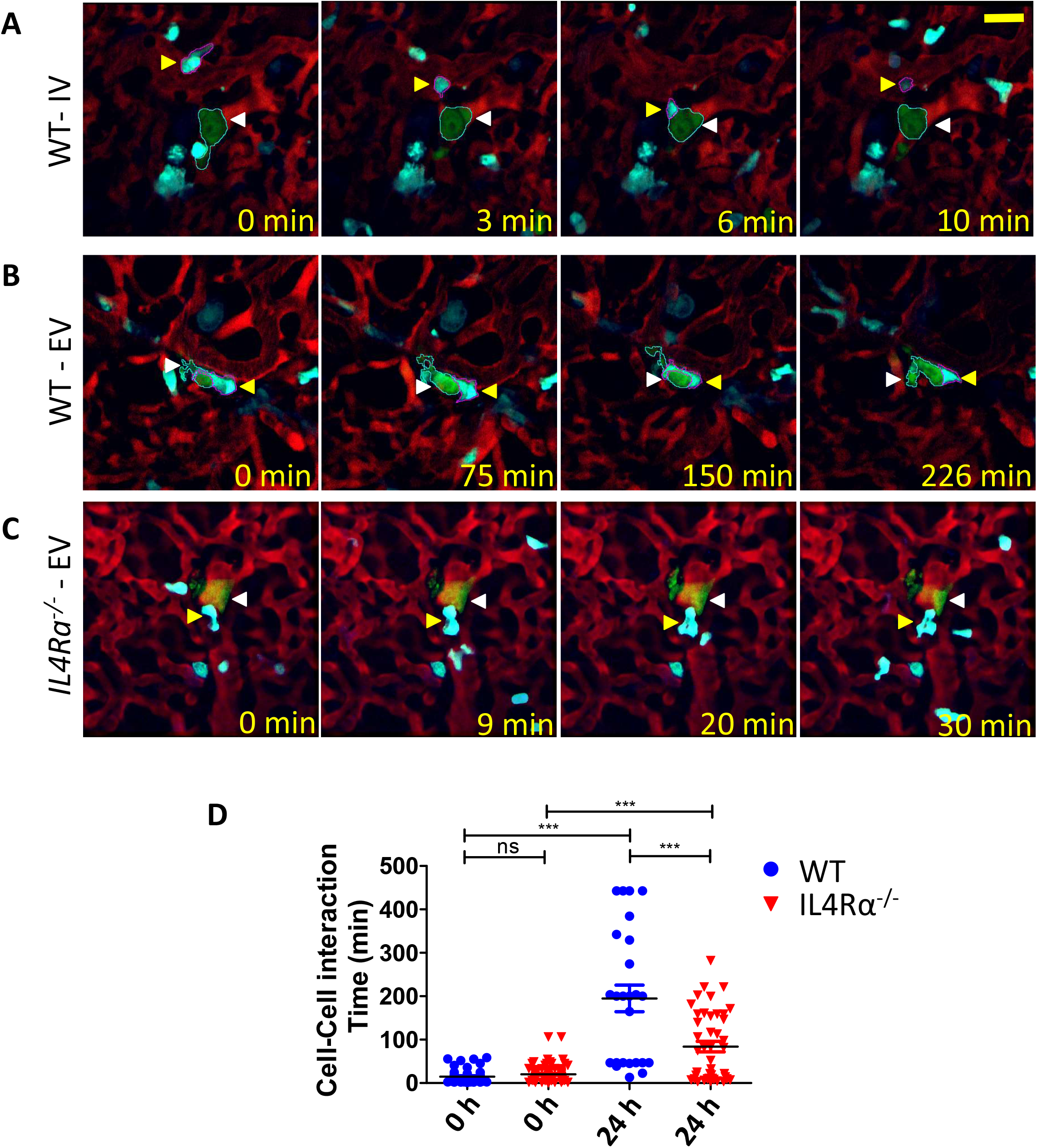
Tumor cell-macrophage interaction in the pulmonary metastatic site *in vivo*. **(A)** Two-photon microscopy of the lungs of WT MacBlue mice, shortly after (0 h post-injection) intravenous injection of Clover^+^ E0771-LG cells. Tumor cell (green, white arrowhead) interacts with CFP^+^ monocyte (cyan, yellow arrowhead) in the intravascular space (vasculature lumen labeled in red by injection of TRITC-Dextran). **(B)** Two-photon microscopy of the lungs of WT MacBlue mice 24 h after intravenous injection of Clover^+^ tumor cells. An extravascular tumor cell (green, white arrowhead) is seen physically interacting with CFP^+^ macrophage (cyan, yellow arrowhead). **(C)** Two-photon microscopy of the lungs of *IL4Rα*^*-/-*^ MacBlue mice 24 h after intravenous injection of Clover^+^ tumor cells. An extravascular tumor cell (green, white arrowhead) is physically interacting with CFP+ cell (cyan, yellow arrowhead). **(D)** Quantification of the time of physical interaction between tumor cells (green) and monocytes (intravascular CFP+ cells) or macrophages (extravascular CFP cells). Note those cells that remain at the base of the X axis have not extravasated. Scale=40um. Mann-Whitney test, n>3-5mice/group, *** p-value<0.0001.

### IL4 *regulates multiple downstream pathways at the transcriptional level in macrophages*

To investigate how IL4 modulates macrophage activity during tumor cell transendothelial migration, downstream transcriptional targets of IL4 signaling in BMDMs were examined. Using public Affimetrix datasets (GEO accession number GSE35435) the gene expression profiles of untreated BMDMs were analyzed using Ingenuity (Core) Pathway Analysis (IPA) and compared to IL4 treated BMDMs. As expected, the top networks and top biological functions altered by IL4 treatment were strongly associated with developmental and immune responses (**Figure 4A**). Among the top canonical signaling pathways affected by IL4 treatment, *IL-8 signaling* was identified (mice lack IL-8 orthologs but the homologues are CXCL1, CXCL2 and CXCL5). Consequently, we investigated the expression of mRNA CXCL1, CXCL2 and CXCL5 in response to IL4 by qRT-PCR. BMDMs stimulated with IL4 highly suppressed mRNA expression of both CXCL1 (n=3; p-value<0.001) and CXCL2 (n=3; p-value<0.0001), whereas CXCL5 did not show significant differences in transcript abundance perhaps because of the variability in response between replicates for this molecule (n=3; p-value=0.40) (**Figure 4B**).

**Figure 4.**
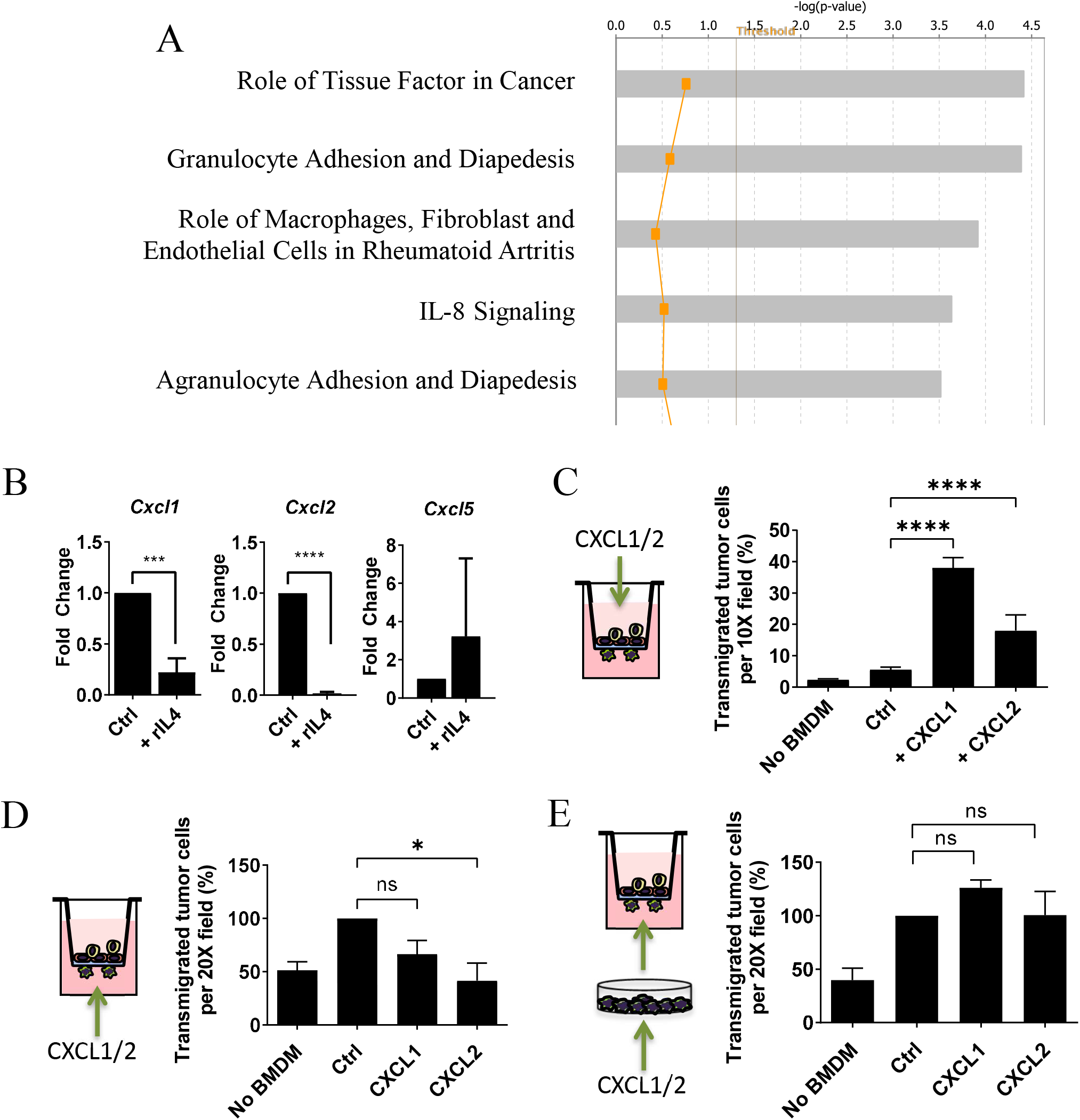
CXCR2 is controlled by IL4 signaling in macrophages and it mediates the IL4 dependent trans-endothelial migration of tumor cells. **(A)** Top 5 canonical pathways derived from ingenuity pathway analysis (IPA) for differentially expressed genes in BMDM stimulated with IL4. These pathways emerged following IPA “Core Analysis.” Graph shows category scores (y axis) based on the p-values derived from the Fisher’s exact test. The x axis displays the –log(p-value). The yellow threshold line represents a significance cutoff at p=0.05. **(B)** Normalized level of mRNA expression measured by qPCR of IL-8 functional homologs CXCL1, CXCL2 and CXCL5 in macrophages after a 24 h stimulation with recombinant IL4 (n=3, *** p-value <0.001; **** p-value<0.0001). **(C)** Number of transmigrated E0771-LG cells in response to WT BMDMs when recombinant CXCL1 or CXCL2 is present at the receiving chamber throughout the assay (n=3, *** p-value<0.005). **(D)** Number of transmigrated E0771-LG cells in response to WT BMDMs in presence of recombinant CXCL1 or CXCL2 at the bottom chamber (n=3, * p-value<0.02). (B-D) Bars represent mean normalized to control +/-SD. (**E**) Number of transmigrated E0771-LG cells in the presence of WT BMDM pre-incubated for 24 h with CXCL1 or CXCL2 (n=3, p-value>0.5).

CXCL1 and CXCL2 are part of a gene-expression signature that mediates experimental metastasis to the lung by primary human breast cancer cells and correlates with larger tumor size in breast cancer patients [34]. Given these data, we hypothesized that CXCL1 and CXCL2 may play a role during tumor cell trans-endothelial migration *in vitro* by establishing chemokine gradients during the eTEM assays. To test this, tumor cell transendothelial migration was stimulated by adding active recombinant chemokines to the top chamber of the transwell insert during eTEM. Under these conditions a significantly higher number of tumor cells transmigrating the endothelial barrier (E0771-LG cells, n=3, p-value<0.02) (**Figure 4C**). To simulate macrophages that also express higher levels of these proteins, we added CXCL1 or CXCL2 to the bottom chamber during eTEM to generate an inverse gradient. This treatment resulted in a significant reduction of E0771-LG trans-endothelial migration but only when CXCL2 was present (n=3, p-value_CXCL1_=0.15 and p-value_CXCL2_=0.02) (**Figure 4D**). In contrast, WT BMDMs pre-incubated with recombinant active CXCL1 or CXCL2 did not induce significant changes in tumor cell trans-endothelial migration compared to untreated BMDMs (E0771-LG cells, n=3, p-value_CXCL1_>0.99 and p-value_CXCL2_=0.86) (**Figure 4E)**. These results suggest that the presence of these chemokine gradients influence tumor cell transendothelial migration.

### CXCR2 is downstream of IL4 in macrophages and mediates the IL4 dependent increase of tumor cell trans-endothelial migration

To investigate whether IL4 stimulation of macrophages modulates transcript abundance of CXCR2 (the receptor for CXCL1 and CXCL2), CXCR2 expression was evaluated in WT BMDMs after treatment with IL4. We observed a significant five-fold increase of the CXCR2 transcript level after exposure to IL4 (n=4; p-value<0.05) (**Figure 5A**). To test the role of CXCR2 signaling in macrophages during tumor cell trans-endothelial migration *in vitro*, CXCR2 signaling was blocked during the eTEM assay. This pharmacological inhibition of CXCR2 was attained using a specific inhibitor (SB332235) which targets CXCR2 but not the closely related receptor CXCR1 [35]. SB332235 effectively inhibited AKT phosphorylation induced by treatment with CXCL1 or CXCL2 in BMDMs (**Supplemental Figure 2C**). SB332235 strongly inhibited tumor cell trans-endothelial migration when added to macrophages at the bottom of the transwell system (**Figure 5B**). CXCR2 inhibition also suppressed the increase in tumor cell trans-endothelial migration in response to CXCL1 (**Figure 5B**). These results suggest that CXCR2 signaling is important for tumor cell trans-endothelial migration. To evaluate whether pharmacological inhibition of CXCR2 signaling is specific to macrophages, the level of CXCR2 expression in all cellular components of the eTEM assay was measured at the protein level by flow cytometry. All cellular components of the eTEM assay, 3B-11 endothelial cells, Met-1 and E0771 tumor cells, as well as BMDMs express CXCR2 (**Figure 5C**), indicating that the reduction of tumor cell trans-endothelial migration observed with pharmacological inhibition of CXCR2 was not necessarily a result of specific inhibition of macrophages. Therefore, to unequivocally evaluate the specific contribution of CXCR2 signaling in macrophages during tumor cell trans-endothelial migration, we isolated BMDMs from *Cxcr2*^-/-^ mice and compared them to WT BMDMs in the eTEM assay. Similar to that observed with *IL4Rα*^*-/-*^ BMDMs, there were no difference in the number of transmigrated tumor cells in response to *Cxcr2*^-/-^ compared to WT BMDMs (Met-1 cells, n=3, p-value=0.92) (**Figure 5D**). Since WT macrophages stimulated with IL4 showed an increased level of *Cxcr2* transcription, it was determined whether increased levels of CXCR2 induced by IL4 mediates the IL4-dependent increased trans-endothelial migration of tumor cells (**Figure 5D**). BMDMs derived from WT and *Cxcr2*^-/-^ mice treated with IL4 were compared using the eTEM assay. IL4 pre-stimulation of *Cxcr2*^-/-^ BMDMs did not increase transmigration of tumor cells (n=3, p-value=0.56), in contrast to the enhanced effect with WT BMDMs (n=3, p-value<0.05) (**Figure 5D**). *CXCR2*^-/-^ BMDM were isolated from a global KO in a BALB/c genetic background. Similar results were observed with E0771-LG cells and *Cxcr2* floxed BMDM from C57BL/6 LysM-Cre mice that deletes genes effectively in macrophages (**Figure 5E**). These data indicate that CXCR2 mediates the IL4 dependent increase of tumor cell trans-endothelial migration of macrophages *in vitro*.

**Figure 5.**
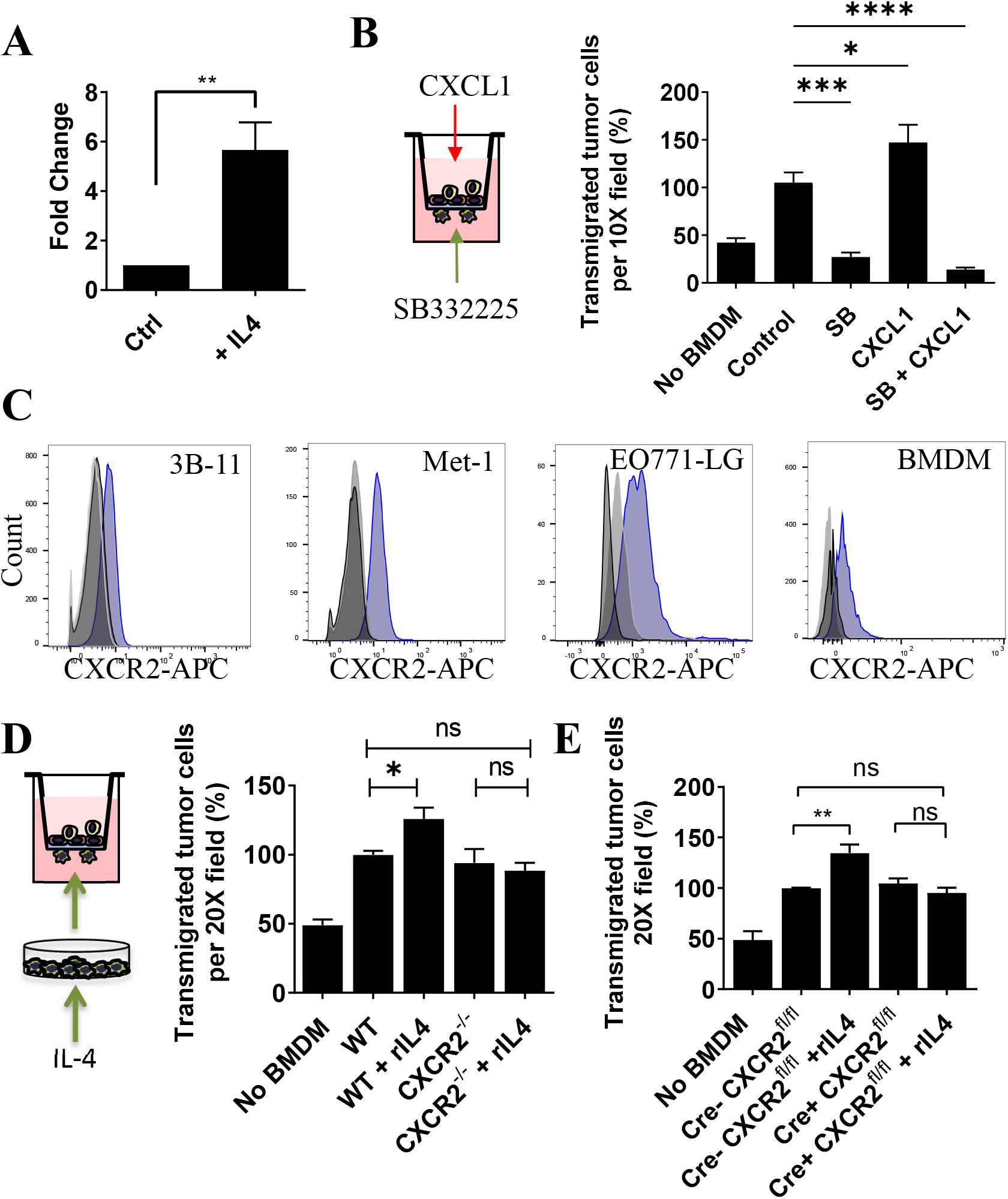
CXCR2 is controlled by IL4 signaling in macrophages and it mediates the IL4 dependent trans-endothelial migration of tumor cells. **(A)** Normalized level of expression of the mRNA *Cxcr2* in macrophages after stimulation with IL4 (n=4, ** p-value <0.01). **(B)** Number of transmigrated E0771-LG cells in response to WT BMDMs treated with 100 μM SB332235 at the bottom chamber, recombinant CXCL1 in the receiving chamber, or both (n=3; **** p-value<0.0001; ** p-value<0.01). **(C)** Surface expression of CXCR2 determined by flow cytometry on 3B-11, Met-1, E0771-LG, and WT BMDM (blue curve) compared to isotype control (light gray) and unstained control (dark gray). **(D)** Transendothelial migration of Met-1 cells in response to WT or CXCR2^-/-^ BMDM isolated from global knock-out Balb/c mice pretreated with recombinant IL4 (n=3; * p-value<0.05). **(E)** Transendothelial migration of E0771-LG cells in response to WT or *Cxcr2*^*fl/fl*^ BMDM with or without Cre expression isolated fromLysM-Cre-CXCR2^fl/fl^ mice (n=4; ** p-value<0.01). For all graphs, bars represent averages normalized to control +/-SD.

### IL4 Regulates genes in macrophages required for metastatic seeding and expansion

Within the gene list related to IL-8 signaling identified by IPA in IL4 stimulated macrophages, *Flt1* was the most upregulated gene. FLT1 is important for the pro-tumoral activities of MAMs in the lung but not for their recruitment [13]. IL4 stimulation of BMDMs induces a significant increase of *Flt1* expression at the transcript level determined by qRT-PCR (n=4, p-value <0.01) (**Figure 6A**). IL4 signaling also induced FLT1 cell surface protein expression **(Figure 6B)**. BMDMs from IL4Rα null mice failed to show an increase FLT1 on their membrane over the base line expression in response to IL4 (**Figure 6B**). Interestingly, other genes associated with macrophage functions during metastatic progression, such as *Ccl2* and *Csf1*, are also controlled at the mRNA and protein level by IL4 signaling (**Figure 6D, E**) [18, 29]. A further qRT-PCR assessment of genes previously shown to be important in lung experimental metastasis assays [18, 36], revealed up-regulation at the transcript level of *Ccr1, Ccl3* and *Hgf* (**Fig 6D**). These data suggest that IL4 signaling can be a master regulator that controls multiple downstream effectors in MAMs that have been shown by genetic ablation to be important during tumor cell metastatic seeding (CCL2, CCR1) and growth (FLT1, CSF1) *in vivo* (**Figure 6F**).

**Figure 6.**
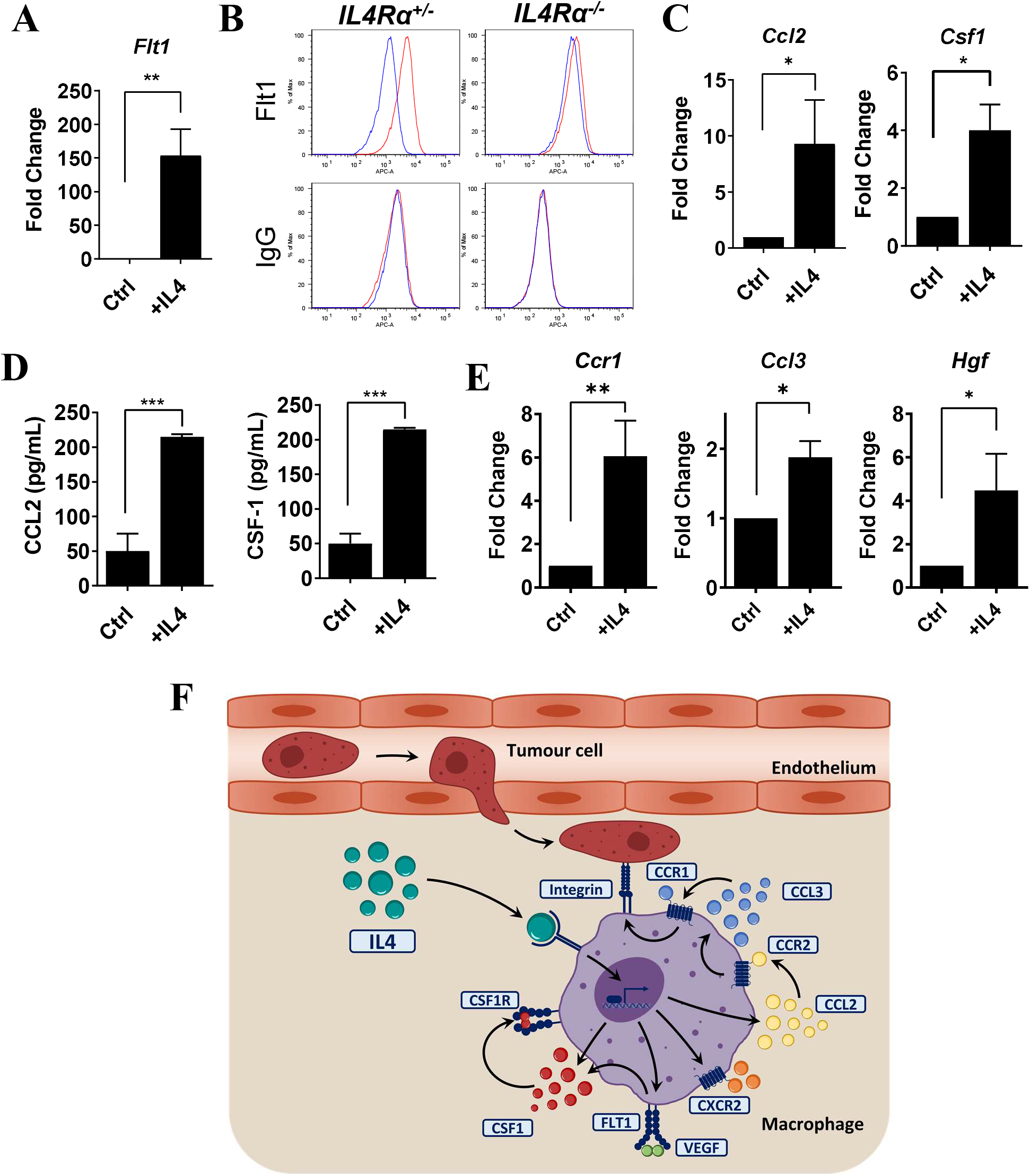
IL4 signaling controls the expression of *Flt1* and other molecules associated with metastatic progression. **(A)** mRNA expression of *Flt1* in BMDM measured by qPCR in response to 24h stimulation with rIL4 (n=4; **p value<0.01). **(B)** FLT expression on *IL4Rα*^+/-^ and *IL4Rα*^-/-^ BMDM cell surface in control conditions (blue line) and after stimulation with rIL4 (red line) measured by flow cytometry. Graph represents the mean fluorescence intensity for each condition. (**C**) mRNA expression of *Ccl2* and *Csf1* in BMDM measured by qPCR in response to 24h stimulation with rIL4 (n=4; *p value<0.05). **(D)** CCL2 and CSF-1 protein expression by BMDM after 24 h of stimulation with rIL4 determined by ELISA (n=3, *** p-value<0.005 and ** p-value<0.01). **(E)** (I) Normalized levels of *Ccr1, Ccl3* <u>and</u> *Hgf* mRNA expression in BMDM from CXCR2^*fl/fl*^ mice after 24 h of stimulation with rIL4, measured by qPCR (n=3, ** p-value<0.01 and *p-value<0.05). (**F**) Proposed model for the action of IL4 effects on macrophages in the metastatic site supported by the data presented in this study and genetic analysis in previous studies.

## Discussion

In the metastatic cascade tumor cells escape into the circulation either directly or via the or lymphatic system and transit to metastatic sites where they extravasate [37]. These sites are often pre-conditioned by systemic influences of cancer causing the creation of the so-called pre-metastatic niche [38]. These niches are often populated by myeloid cells [39]. Tumor cells can travel either singly or in clumps but in either case the rate limiting steps is in their survival and subsequent establishment as metastatic lesions [5, 40]. In human breast cancer the major sites of metastasis are bone and lung, and these are the major causes of mortality in this disease accounting for ∼90% of deaths [41]. To study these diseases, we and others have developed experimental and spontaneous mouse models of lung and bone metastases [5]. In these models of metastasis, once the tumor cells arrive, there is an immediate CCL2 mediated recruitment of CCR2 expressing classical monocytes that promote metastasis seeding and establishment [16]. In lung, these classical monocytes differentiate via a progenitor state (MAMPC) into mature MAMs, each step promoting a different aspect of tumor cell growth into a metastatic lesion [18]. Thus, removal of these cells by blocking their recruitment, function or survival inhibits metastasis [16]. In this study we show that IL4, classified as a Th2 cytokine, regulates the function of these cells through type 1 IL4 receptor signaling. This signaling induces a gene expression cascade in monocytes as they differentiate into MAMs that includes the upregulation of *Flt1, Csf1, Cxcr1, Hgf* and *Ccr1*. IL4 is therefore a master regulator of monocytes and MAMs causing them to promote the seeding, survival, and persistent growth of metastatic cells.

During cancer evolution it has been suggested that the tumor microenvironment needs to become immunosuppressive, so that tumors escape immune responses and thus flourish [14, 42]. Part of the evidence for this immunoregulation was shown with allografted tumors that in some contexts thrived in Th2 biased (Balbc) but were rejected in Th1 mouse strains (BL6) [43, 44]. In this case rejection was mediated by iNOS expressing macrophages in the BL6 mice while in the Balbc strain tumor growth was promoted by macrophage derived arginase that is immunosuppressive. Consequently, the macrophages in these tumors from these two strains were labelled as M1 or M2 to match the Th1/2 nomenclature [43]. In tissue culture, Th1 cytokines, IL12, IL18, INFγ and Th2 cytokines, IL4, IL10 and IL13, cause macrophages to differentiate to an activated or alternatively activated state respectively. These states are characteristic of in vivo responses to bacterial or helminth infections respectively [45, 46]. Thus, the original M1 to M2 definition was extended to encompass these activated (M1) and alternatively activated (M2) states with the consequent suggestion that M2s are tumor promoting [47-49]. Subsequent research in mouse models of cancer indicated in primary tumors that IL4 is an essential cytokine responsible for macrophage differentiation to support a state that promotes macrophage enhanced tumor cell invasion and thus tumor malignancy [21, 50].

While the function of IL4 in the primary tumor has been well studied, its role in metastasis particularly those to the lung has not been explored. In the current study using models of breast cancer metastasis to the lung, we indicate that IL4 regulates aspects of the differentiation of monocytes to MAMs. Consistent with this action ablation of IL4 signaling inhibited metastatic cell seeding, survival and persistent growth. Furthermore, IL4 stimulates the expression of several downstream genes that in previous experiments have been shown to be essential for this process (**Figure 6F**). This signaling include expression of CCR1 required for MAM adhesion to extravasated tumor cells that in turn delivers a survival signal to these cells [18]. IL4 also stimulates FLT1 (VEGFR1) cell surface expression that makes the MAMs responsive to VEGFA and CSF1 that acts in an autocrine fashion through the CSF1R and stimulates macrophage survival and polarization [29]. During these processes MAMs differentiate via a precursor that is characterized by a Th2 gene expression signature and is immunosuppressive to cytotoxic T cells through superoxide formation [51]. These cells are often referred to as myeloid derived suppressor cells. Upon their differentiation to MAMs, driven by CSF1R and FLT1 signaling, they obtain a new suppressive activity to cytotoxic T cell killing via expression of CTLA-4 check-point inhibitor ligands [51]. In addition, IL4 induces expression of HGF that creates an immunosuppressive environment to cytotoxic Natural Killer cells via c-MET signaling in tumor cells [36]. Thus in this model of breast cancer lung metastasis, IL4 generates a immunosuppressive Th2 environment through its actions on monocyte differentiation to MAMs after their recruitment via CCL2 [16].

Monocytes and MAMs in addition to creating an immunosuppressive environment, also promote the earliest steps in metastasis by enhancing metastatic cell extravasation. This action is in part due to the production of VEGFA that increases vascular permeability [16]. The eTEM assay in our hands has been valuable to analyze this step of extravasation. This study showed that IL4 is not required for macrophage enhancement of the transendothelial migration of tumor cells but does enhance the macrophage’s ability to promote this process. IL4 upregulates CXCR2 whose genetic deletion specifically in macrophages inhibits their ability to promote this process. IL4 also downregulated the ligands CXCL1 and 2 suggesting that this removal of an autocrine response might promote paracrine signaling through CXCR2. In fact, pre-incubation of the BMMs with CXCL1 inhibited their ability to enhance transendothelial migration, suggesting the downregulation of CXCL1 might allow interaction of CXCR2 with another ligand such as, CXCL8, that has been shown to enhance breast cancer metastasis through action on myeloid cells [52].

Importantly in the studies reported here, we also used a novel quantitative intravital imaging method to analyze interactions between tumor cells and monocytes/MAMs during the stages of intravasation *in vivo*. These data unequivocally showed that IL4 signaling enhanced tumor cell/MAM interactions and stabilized the association between these two cell types. Previous studies indicated that this association was through the tumor cell surface expression of VCAM1 and alphaV integrin on the MAMs whose activity was stimulated by CCR1 signaling [18]. This engagement delivers a survival signal to the tumor cells and thereby promotes metastatic cell colonization [18, 53]. It also promotes monocyte/MAM retention at the metastatic site [18]. IL4 stimulates this process at least in part, by upregulation of CCR1 and CCL2 expression. CCL2 in turn, upregulates CCL3 expression that stimulates CCR1 activity [53]. These real time *in vivo* experiments therefore validate the activities found *in vitro* [18] and indicate that IL4 is a major regulator of the process as summarized in Figure 6F.

While the experimental metastasis methodology used in this study has its limitation because cells highly selected for colonization into the bone or lung are given as a bolus of single cells in high concentration, it does allow dissection of the process without influences of the primary tumor that can confound interpretation. However, consistent with the results obtained in the experimental metastasis model, null or conditional mutations or specific inhibitors (antibodies/small molecules) in almost all the genes described in this study, including the *IL4Ra, Flt1, Vegfa* and *Csf/Csf1r* inhibits spontaneous metastasis in mouse models of breast cancer [18, 21, 29]. In models of breast cancer bone metastasis, a tissue where autochthonous tumors do not metastasize to in mice, classical monocytes recruitment has also been shown to be essential for metastatic progression [27]. These monocytes differentiate into Bone MAMs (BoMAMs) that upregulate IL4Ra and whose signaling promotes metastatic outgrowth [27]. Furthermore, IL4R positive BoMAMs have been shown in osteolytic lesions caused by human breast cancer metastasis [27]. Together these data suggest that IL4R inhibition might be a therapeutic avenue to inhibit Th2 suppression of cytotoxic immune response and enhance immunotherapy for deadly bone and lung metastatic disease.

## Methods

*Animal models*. All experiments using animals were performed in accordance with protocols approved by the Albert Einstein College of Medicine Institutional Animal Care and Use Committee (IACUC). Femurs and tibias from *Stat3*^*fl/fl*^ (*Tg(Csf1r-iCre)jwp*^*-/-*^*Stat3*^*flox/flox*^) and *Csf1r*.*iCre;Stat3*^*fl/fl*^ (*Tg(Csf1r-iCre)jwp*^*+/-*^*Stat3*^*flox/flox*^) were donated by Dr. Elaine Lin (Montefiore Medical Center, Bronx, NY) [30]. B6-*IL4rα*^*tm1Sz*^/J (*IL4Rα-/-*) mice were donated by Dr. Johanna Joyce (Memorial Sloan Kettering Cancer Center, NY, NY). Mice carrying CXCR2 homozygous deletion (C.129S2(B6)-*Cxcr2*^*tm1Mwm*^/J) and their WT (BALB/cJ) control were purchased from Jackson Laboratories. Femurs and tibias from animals carrying the tissue specific CXCR2 homozygous deletion in myeloid cells (LysM-Cre recombinase x C57BL/6-*Cxcr2*^*tm1Rmra*^) and their Cre-controls were kindly donated by Dr. Ann Richmond (Vanderbilt University). MacBlue (B6,FVB-Tg(Csf1r-GAL4/VP16,UAS-ECFP)1Hume/J) mice [54] were purchased from Jackson Laboratories. WT FVB/NJ mice were also purchased from Jackson laboratories. All animals were housed in SPF conditions.

### Cell lines and tissue culture

Mouse E0771 medullary mammary adenocarcinoma cells were obtained from Dr. E. Mihich (Rosewell Park Cancer Institute, New York). E0771 cells were originally isolated from a spontaneous mammary tumor in C57BL/6 mice [55]. Highly metastatic lung-tropic E0771 tumor cells (E0771-LG) were previously derived from the parental cell line by isolation from lung foci after repeated cycles of experimental metastasis assays [18]. Murine endothelial 3B-11 cells were obtained from the ATCC^R^ (CRL-2160™). Met-1 cells were derived from a spontaneous mammary carcinoma from the MMTV-PyMT mouse model in FVB mice [56]. All cell lines were maintained in Dulbecco’s modified Eagle medium (DMEM) (Invitrogen) supplemented with GlutaMAX™-I, 10% v/v heat-inactivated fetal bovine serum (FBS), 1mM sodium pyruvate, 100U/mL of penicillin, and 100 μg/mL of streptomycin, at 37°C and 5% v/v CO_2_ (complete DMEM). The fluorescent E0771-LG cell line expressing Clover (Addgene) [57] used in intravital experiments was established by standard transfection, as previously published. To obtain these, E0771-LG cells were cultured overnight in 10% v/v FBS-DMEM without antibiotics and transfected with lipofectamine® 2000 together with the CLOVER plasmid. Cells were cultured in selection media (700 μg/mL G418 in complete DMEM) for a week and enriched for the 10% brightest population using fluorescence-activated cell sorting (FACS). Sorted cells were maintained under selection for a second week and enriched once again for the 10% brightest population by FACS. The cell line established was evaluated for compatibility with intravital imaging (brightness, photobleaching and photodamage). E0771-LG cells expressing Clover were subjected to a third round of selection in which single cell clones were isolated into 96 well plates by FACS. All intravital imaging sessions were performed using two clones, #14 and #15.

### Primary culture of bone marrow derived macrophages

Femurs and tibias from 8-12 week-old mice were collected and flushed with a 23-G needle in cold Alpha Minimum Essential (αMEM) Medium. The bone marrow cell suspension was centrifuged at 180 g for 5 minutes and plated overnight in tissue culture plates in 10% v/v heat inactivated FBS, 100U/mL penicillin, 100 μg/mL streptomycin, and 10^4^ U/mL of human recombinant (rh) CSF1 (complete αMEM), at 37°C and 5% v/v CO_2_. Non-adherent cells were then transfered to petri dishes and cultured in complete αMEM for 6 days. Cells were supplemented with fresh complete αMEM every 3 days and used for experiments no later than 2 weeks after bone marrow isolation.

### Endothelial cell permeability assay and trans-endothelial resistance

Matrigel(R)-coated transwell inserts (Corning) with 8 μm pores were thawed at room temperature for 10 min and pre-incubated with complete DMEM in both chambers for two hours at 37°C. 10^4^ 3B-11 cells were seeded on top of the filter in complete DMEM at different time points. For trans-endothelial resistance (TER) assays, medium was replaced with DMEM containing 0.5% v/v FBS, 1mM sodium pyruvate, 100 U/mL Penicillin, and 100 μg/mL Streptomycin (0.5% v/v FBS-DMEM) two hours before measurement. Trans-endothelial resistance was measured using EndOhm-6 chamber and EVOM^2^ resistance meter (World Precision Instruments) and expressed as a function of the effective surface area of the filter membrane (Ω x cm^2^). For permeability assays, media in the receiving (top) chambers was replaced with DMEM containing 0.5% v/v FBS, 1mM sodium pyruvate, 100 U/mL Penicillin, 100 μg/mL Streptomycin, and 100 μg/mL of 70 kD Rhodamine B Isothiocyanate-Dextran (TRITC) and incubated for 2 hours. Aliquots of 100 μL from the receiving and bottom chambers were taken and diluted 1:5 in DMEM before reading using a SpectraMax M5 plate reader (emission/excitation = 557/586 nm; Molecular Devices). Concentrations were calculated using a standard curve. Control condition consists of transwell inserts with no 3B-11 cells, incubated for 2 hours in 0.5% v/v FBS-DMEM.

### Experimental trans-endothelial migration assay

Matrigel(R)-coated transwell inserts (Corning) with 8 μm pores were thawed at room temperature for 10 min and pre-incubated with complete DMEM in both chambers for two hours at 37°C. 10^4^ 3B-11 murine endothelial cells were seeded in 200 μL of complete DMEM in the receiving chamber (top), with 750 μL of complete DMEM in the bottom chamber. Cells were incubated for 48 h at 37°C and 5% v/v CO_2_ until confluent. E0771-LG cells and Met-1 cells were labeled with CellTracker™ Green CMFDA Dye (Thermo Fisher Scientific, New York) following the manufacturer’s instructions, trypsinized and adjusted to 1 × 10^5^ cells/mL (Met-1 cells) or 2 × 10^5^ cells/mL (E0771-LG cells) in 0.5% v/v FBS-DMEM containing 10^4^ U/mL hrCSF1. Bone marrow-derived macrophages (BMDMs) were lifted into a cell suspension with a rubber policeman and adjusted to 5×10^5^ cells/mL in complete DMEM supplemented with 10^4^ U/mL of rhCSF1. Macrophages were seeded on the bottom side of the transwell insert by carefully depositing 20 μL of the cell suspension and allowing attachment for 30 min at room temperature. Transwell inserts were moved to wells containing 750 μL of complete DMEM with 10^4^ U/mL of rhCSF1 and received 200 μL of tumor cell suspension in the top chamber. After 36 h, cells at the receiving chamber were removed using a cotton swab. Cells at the bottom chamber were fixed in 4% w/v paraformaldehyde (PFA) in PBS for 30 min at 4°C. Six randomly selected fields were acquired per transwell insert in an inverted Olympus IX81 microscope equipped with a 20X (NA=0.40 Air) objective and a Sensicam QE cooled CCD camera (Analytical Imaging Facility, Albert Einstein College of Medicine). Images were quantified using Fiji / ImageJ software [58]. Each experiment was performed in a minimum of three independent repeats, and in triplicate. BMDMs, unless otherwise specified, were stimulated with cytokines for 24 h at 10 ng/mL. Recombinant murine IL4, IL13, IL10, CXCL1, CXCL2 and CXCL5 were purchased from Peprotech and reconstituted in sterile 0.05% w/v BSA-PBS and stored at -80°C. Blocking antibody against IL10R was used at 20ng/mL (Rat anti-mouse CD210, BD#550012). CXCR2 inhibitor SB332235 (GlaxoSmith) was diluted in DMSO and used at 100 μM in culture medium.

### Experimental Metastasis Assay

Female mice of 8-12 weeks of age were intravenously injected with 6×10^5^ syngeneic tumor cells through the tail vein in a total volume of 200 μL of PBS as described [18]. Injected mice were maintained under quarantine until tissue collection. Mice were euthanized by isofluorane overdose and lungs were perfused with 5-10 mL of PBS injected to the right ventricle of the heart. Lungs were filled with 1mL of 4% w/v PFA in PBS and fixed overnight in the same fixative. Metastatic burden was assessed by serial sectioning of PFA-fixed paraffin-embedded lung tissue sections stained with hematoxylin and eosin (H&E) using the stereological method [59] or surface nodule count.

### Adoptive Transfer Experiments

Donor female mice of 8-12 weeks of age were euthanized by isofluorane overdose and femurs, tibias, ileac bone, and spine were collected to isolate bone marrow cells. Bones were crushed using mortar and pestle in sterile FACS buffer (1% w/v BSA in PBS) and filtered through a 40 μm cell strainer to obtain single cell suspensions. Cell suspensions were centrifuged at 280 g and 4°C for 5 min and erythrocytes were eliminated by incubation in RBC buffer for 10 min on ice. Cells were counted and adjusted to 2.8 ×10^8^ cells/mL in MACS buffer (1x PBS, 0.5% v/v FBS, 2mM EDTA). Monocytes were enriched by negative selection using a Monocyte Isolation Kit (Miltenyi Biotech, Kit #130-100-629) following the manufacturer’s instructions. After magnetic separation, inflammatory monocytes (CD45^+^CD11b^+^Ly6G^-^Ly6C^+^) were collected using a FACS Aria unit (Becton Dickinson) in sterile conditions. Inflammatory monocytes and tumor cells were injected in a ratio 1:1 through the tail vein.

### Real-time PCR

Total RNA was isolated from cells by homogenization in TRIzol (Thermo Fisher) or RNeasy miniprep kit (Qiagen) and converted into cDNA by using SuperScript® Vilo (Thermo Fisher). Gene expression was determined by real-time PCR using SyBRGreen (Invitrogen) and showed as relative gene expression after normalization to *GAPDH*. The following primers were used: *Gapdh* forward: 5’-CTTTGTCAAGCTCATTTCCTGG-3’; *Gapdh* reverse: 5’-TCTTGCTCAGTGTCCTTGC-3’; *Cxcl2* forward: 5’-AATGCCTGAAGACCCTGC-3’; *Cxcl2* reverse: 5’-TTTTGACCGCCCTTGAGAG-3’; *Ccl3* forward: 5’-ATGAAGGTCTCCACCACTGC-3’; *Ccl3* reverse: 5’-CCCAGGTCTCTTTGGAGTCA-3’; *Ccl5* forward: 5’-GTTCCATCTCGCCATTCATG-3’; *Ccl5* reverse: 5’-TTAAGCAAACACAACGCAGC-3’; *Cxcl1* forward: 5’-AGAACATCCAGAGCTTGAAGG-3’; *Cxcl1* reverse: 5’-CAATTTTCTGAACCAAGGGAGC-3’; *Cxcr2* forward: 5’-CGTTTGAGGGTCGTACTGCG-3’; *Cxcr2* reverse: 5’-TGGGCCTTAAAGAGGGTGCG-3’; *Flt1* forward: 5’-GAGCCAGGAACATATACACAGG-3’; *Flt1* reverse: 5’-GCTTGACAGTCTAAGGTCGTAG-3’; *Vegf1* forward: 5’-AAAGCCAGCACATAGGAGAG-3’; *Vegf1* reverse: 5’-ATTTAAACCGGGATTTCTTGCG-3’; *Il13* forward: 5’-CCAGCCCACAGTTCTACAG-3’; *Il13* reverse: 5’-GAGACACAGATCTTGGCACC-3’.

### Ingenuity Pathway Analysis

Datasets comparing BMDMs treated with recombinant murine IL4 for 24 h at 20ng/mL and untreated control conditions were accessed from the NCBI database. Gene list with accession number GSE35435 [60] was uploaded to Ingenuity Pathway Analysis tool (Qiagen) and curated for differentially expressed genes (p-value<0.05) and fold change>1.5.

### Flow cytometry

Blood samples were collected through cardiac puncture or retro-orbital bleeding in heparin. Single cell suspensions were incubated for 10 min in red blood cell (RBC) lysis buffer (Biolegend) on ice to eliminate erythrocytes. Cell suspensions were incubated for 30 min on ice with anti-mouse Fc Block CD16/32 antibody (eBiosciences) in FACS buffer to avoid non-specific antibody binding. Cells were then washed in FACS buffer and stained with either Ig controls or fluorophore conjugated antibodies in FACS buffer. Data acquisition was performed on a LSR II Yellow (BD Biosciences) and analyzed on FlowJo version 10 (FlowJo, LLC). Classical (CM: CD45^+^ CD11b^+^ F4/80^+^ Ly6G^-^ Ly6C^hi^) or non-classical patrolling monocytes (PM: CD45^+^ CD11b^+^ F4/80^+^ Ly6G^-^ Ly6C^-^) were identified by FACS using the above antibodies as described [16]. Antibodies used were: anti-CD45-PerCpCy5.5 (eBioscience #45-0451-82); CD11b-eVolve 605 (eBioscience #83-0112-42); F4/80-FITC (Biolegend #123108); Ly6C-PE-Cy7 (Biolegend #128018); Ly6G-APC-Cy7 (Biolegend #127624); CD115-APC (eBioscience #17-1152-82); CD11b-PECy7 (Biolegend #101216); Ly6C-APC-Cy7 (BD-560596); Ly6G-PE (BD#551461).

### Intravital Imaging

Female WT or *IL4rα*^-/-^ MacBlue mice of 6-12 weeks of age were prepared for surgery as previously described [31, 32]. Mice were connected to a mechanical ventilator (Harvard Apparatus, Massachusetts) providing 0.2 cc of air at ∼140 cycles per second and 3% v/v of isofluorane. Then, skin and muscle layers of the upper left thorax were removed. The largest lung lobule was exposed by resection of ribs and intercostal muscles. A vacuum-stabilized lung window [58] was attached to the exposed lung area. Mice were maintained under light anesthesia at physiological temperature and hydrated by periodic injections of saline. Vital signs were monitored using a pulse oximeter (MouseOx, Starr LifeSciences). At the end of imaging sessions, animals were euthanized by isofluorane overdose and cervical dislocation. Vasculature was labeled with an intravenous injection of 155kD TRITC Dextran (Sigma-Aldrich). Imaging sessions were performed with intravenous delivery of up to 2×10^6^ Clover expressing E0771-LG cells in *IL4rα*^+/+^ MacBlue or *IL4rα*^-/-^MacBlue mice. Imaging was performed on a custom made 2-laser-2-photon microscope (Gruss Lipper Biophotonics Center, Albert Einstein College of Medicine) [58]. Multiple positions were imaged from 3 to 12 hours in a 4D format (z-stack + time). Images were analyzed using Fiji/ ImageJ. At the end of each imaging session, animals were euthanized by overdose of isofluorane and cervical dislocation.

### Statistical Analysis

Comparisons between two groups were evaluated by Student’s unpaired t-test. Comparisons between multiple groups were evaluated by one-way analysis of variance (ANOVA) with Dunnet correction for multiple comparisons. Comparisons between groups for *in vivo* experiments were evaluated by Mann-Whitney or Kruskal-Wallis test. Adjusted p-values are shown, unless otherwise indicated. All statistical analyses were performed using the Prism-GraphPad software.

## Author Contributions

CRT performed the majority of experiments and data analysis, prepared figures and wrote the manuscript. CRT and DE performed intravital imaging and analysis with advice from JSC; JL performed WB and FC experiments; BQ performed FC experiments, JWP supervised experiments, data analysis and interpretation, and wrote the manuscript.

## Acknowledgments

We thanks Dr. Joanna Joyce for sharing the C57BL/6 IL4rα null mice, Dr. Amit Verma for sharing the CXCR2 inhibitor SB332285, Dr. Fiona Yull for sharing bone marrow from Stat3 null animals, Dr. Ann Richmond for sharing bone marrow from CXCR2 null mice, and Einstein’s Analytical Imaging Facility (AIF) for imaging support. We thanks Dr Nikki Graham for drawing the summary figure.

This research was supported by the Wellcome Trust 101067/Z/13/Z (JWP), MRC Centre grant MR/N022556/1 (JWP), The Gruss-Lipper Biophotonics Center and its Integrated Imaging Program (DE, JSC), Jane A. and Myles P. Dempsey (JSC), NIH grants P01 CA100324 (JWP/JC), R01CA131270 (JWP), P30 CA013330 (AIF), and R01 CA216248 (DE, JSC).

---

**Supplementary figure 1.**
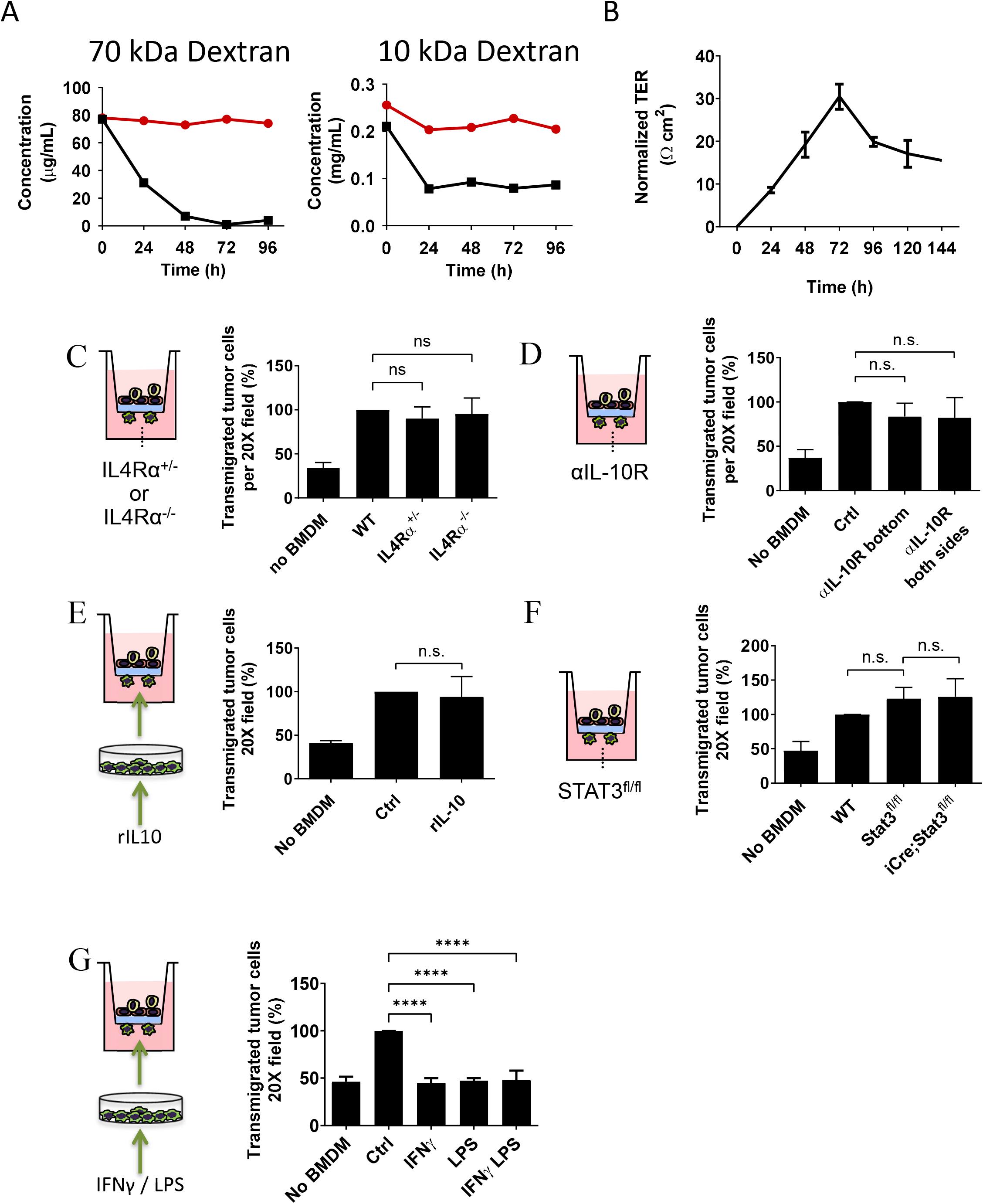
*In vitro* Extravasation assay. (**A**) Permeability assay for 3B-11 endothelial cells cultured on Matrigel-coated transwell inserts with 8 μm pores. Permeability to apically delivered Rhodamine B Isothiocyanate of 2 molecular weights was determined at different times by measuring presence of the fluorescent dye at the receiving (red line) and bottom (black line) side of the transwell. (**B**) 3B-11 cells cultured in the same conditions as in (A) and trans-endothelial resistance (TER) was determined. Highest TER values (n=6, p-value<0.0001) coincide with lower permeability rates, period of time when eTEM assays were performed. (**C**) Transendothelial migration of E0771-LG cells was measured in response to WT, *IL4Rα*^*+/-*^ or *IL4Rα*^*-/-*^ BMDMs. Mann-Whitney test, n=4, p-value>0.8. **(D)** Number of transmigrated E0771-LG cells in response to WT BMDMs in presence of IL10R blocking antibody at the bottom or both chambers (n=3, p-value>0.64). **(E)** Number of E0771-LG transmigrated cells in presence of WT BMDM pre-incubated with 10ng/mL of recombinant IL10 (n=3, p-value=0.97). **(F)** Number of transmigrated E0771-LG cells in the presence of WT, *STAT3*^*fl/fl*^ and iCre^+^*STAT3*^*fl/fl*^ BMDMs (n=4, p-value>0.1). **(G)** Number of transmigrated E0771LG cells in the presence of WT BMDM pre-incubated with recombinant IFNγ, LPS or a combination of both (unpaired one-way ANOVA, n=5, p-value>0.0001). Data are shown as mean normalized to WT or untreated control +/-SD.

**Supplementary figure 2.**
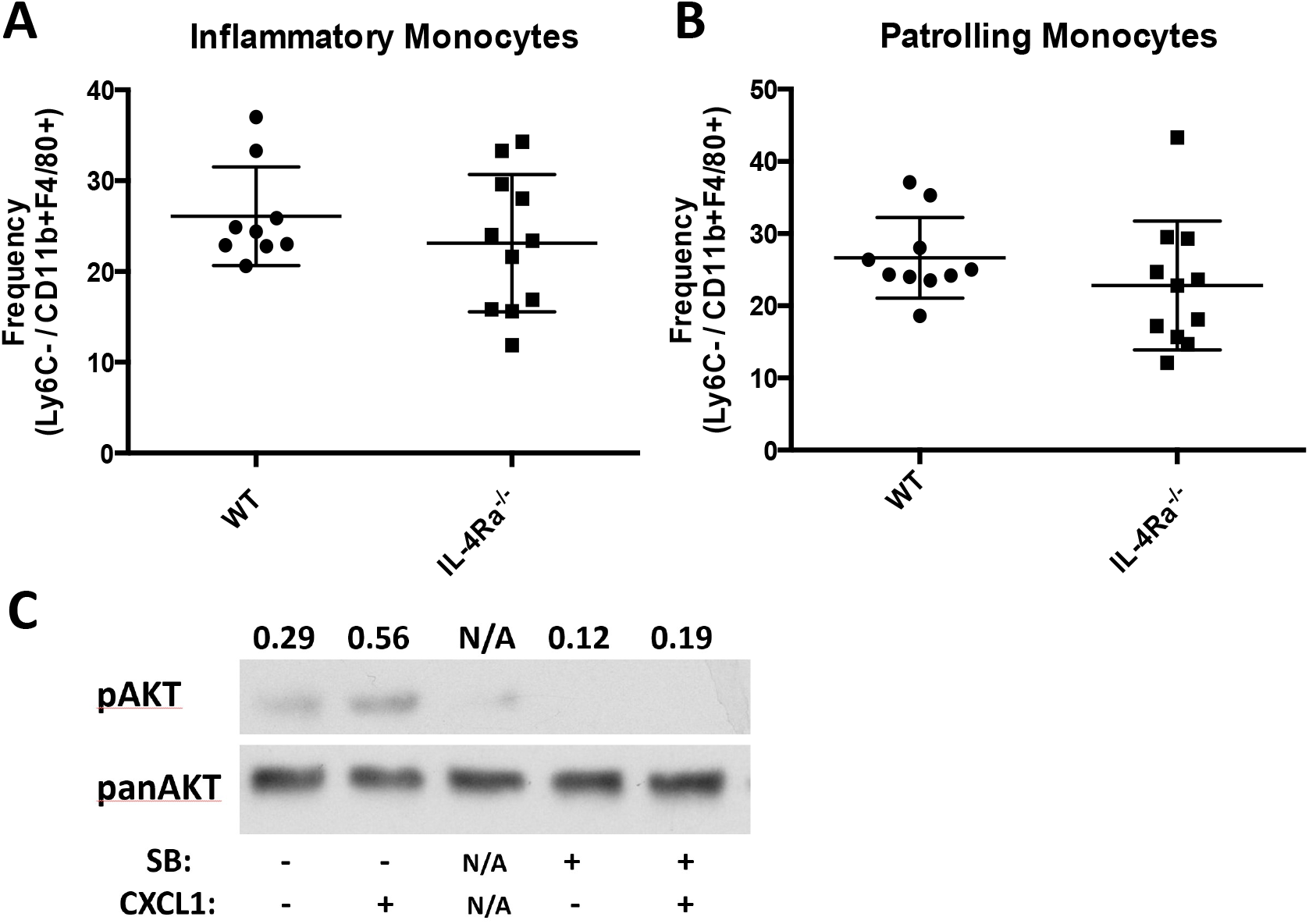
IL4Rα null mice have normal numbers of classical and non-classical monocytes. **(A)** Frequency of classical monocytes (CD45^+^CD11b^+^F4/80^+^Ly6^Hi^) respect to parental population (Live CD45^+^CD11b^+^F4/80^+^) in WT and IL4Rα^*-/-*^ null mice (n>9, p-value>0.1). (**B**) Frequency of non-classical monocytes (CD45^+^CD11b^+^F4/80^+^Ly6^-^) respect to parental population (Live CD45^+^CD11b^+^F4/80^+^) in WT and IL4Rα^-/-^ null mice (n>9, p-value>0.1). (**C**) WB for phosphoAKT and total (pan)AKT in BMDM stimulated with 20 ng/mL CXCL1 alone or in combination with 100 μM SB332235 (SB) for 24 h. Unrelated sample labeled as N/A. Densitometry analysis for the ratio of pAKT/panAKT is shown.

**Supplementary Movie 1**. EO771-LG-Clover cells (green) were intravenously injected into IL4Rα^+/+^ MacBlue mice and immediately after injection, intravital imaging of the lungs was carried out on a custom made 2-laser-2-photon microscope for 3-12 hrs. Movie represents a total time of 10 min. Movie is one representative region from one mouse. Monocytes (intravascular) and macrophages (extravascular) are labeled in cyan after expression of CFP in MacBlue mice. Blood vessels labeled in red by injection with 155kD TRITC Dextran. Movie correspond to still images from **Figure 3A**.

**Supplementary movie 2**. Intravital imaging of the lungs of live IL4Rα^+/+^ MacBlue mice after 24 h of intravenous injection with syngeneic EO771-LG-Clover cells. Blood vessels labeled in red by injection with 155kD TRITC Dextran. Movie represents a total time of 226 min (∼4h). Movie is one representative region from one mouse. Movie corresponds to still images from **Figure 3B**.

**Supplementary movie 3**. Intravital imaging of the lungs of live IL4Rα^-/-^ MacBlue mice after 24 h of intravenous injection with syngeneic EO771-LG-Clover cells. Blood vessels labeled in red by injection with 155kD TRITC Dextran. Movie represents a total time of 30 min (0.5 h). Movie is one representative region from one mouse. Movie corresponds to still images from **Figure 3C**.

## Notes

JWP is a co-founder, board member and consultant to Macomics LTD an immuno-oncology company. The other authors declare no potential conflicts of interest

